# A missense variant in *CREBRF* is associated with taller stature in Samoans

**DOI:** 10.1101/690586

**Authors:** Jenna C. Carlson, Samantha L. Rosenthal, Emily M. Russell, Nicola L. Hawley, Guangyun Sun, Hong Cheng, Take Naseri, Muagututi‘a Sefuiva Reupena, John Tuitele, Ranjan Deka, Stephen T. McGarvey, Daniel E. Weeks, Ryan L. Minster

**Affiliations:** Department of Human Genetics, Graduate School of Public Health, University of Pittsburgh, Pittsburgh, PA, USA; Department of Biostatistics, Graduate School of Public Health, University of Pittsburgh, Pittsburgh, PA, USA; Department of Chronic Disease Epidemiology, School of Public Health, Yale University, New Haven, CT, USA; Department of Environmental Health, College of Medicine, University of Cincinnati, Cincinnati, OH, USA; Ministry of Health, Government of Samoa, Apia, Samoa; Lutia i Puava ae Mapu i Fagalele, Apia, Samoa; Department of Public Health, Government of American Samoa, Pago Pago, American Samoa, USA; International Health Institute and Department of Epidemiology, School of Public Health, Brown University, Providence, RI, USA; Department of Anthropology, Brown University, Providence, RI, USA

**Keywords:** anthropometry, genetic association, diverse populations, height

## Abstract

**Objectives:** Studies have demonstrated that rs373863828, a missense mutation in *CREBRF*, is associated with a number of anthropometric traits including body mass index (BMI), obesity, percent body fat, hip circumference, and abdominal circumference. Given the biological relationship between height and adiposity, we hypothesized that the effect of this variant on BMI might be due in part to a previously untested association of this variant with height.

**Methods:** We tested the hypothesis that minor allele of rs373863828 is associated with height in a Samoan population in two adult cohorts and in a separate cohort of children (age 5 - 18 years old) using linear mixed modeling.

**Results:** We found evidence of a strong relationship between rs373863828 and greater mean height in Samoan adults (0.77 cm greater average height for each copy of the minor allele) with the same direction of effect in Samoan children.

**Conclusions:** These results suggest that the missense variant rs373863828 in *CREBRF*, first identified through an association with larger BMI, may be related to an underlying biological mechanism affecting overall body size including stature.

## Introduction

Recently, we identified a novel association between a missense mutation (rs373863828, c.1370G>A, p.R457Q) in the gene for transcription factor *CREB3 regulatory factor* (*CREBRF*) and body mass index (BMI) in individuals of Polynesian ancestry (Krishnan et al., 2018; Minster et al., 2016) Despite fine-mapping and functional work suggesting the causal variant of this association is rs373863828 (Minster et al., 2016), the biological mechanism through which *CREBRF* operates to affect BMI is still unknown.

The rs373863828 variant is common in Samoans (minor allele frequency, MAF=0.259), but is very rarely seen in individuals without Pacific Island ancestry (gnomAD MAF for all populations=3.53 × 10^−5^; accessed 12 April 2019). However, this variant is polymorphic and associated with BMI in other Pacific Island populations including New Zealand Māori, Pukapukans, and other Polynesians (Krishnan et al., 2018).

This missense variant is associated with a constellation of anthropometric traits including obesity, percent body fat, hip circumference, and abdominal circumference (Minster et al., 2016). Of those traits, rs373863828 was most significantly associated with BMI, with an estimated 1.36–1.45 kg/m^2^ (*p*_meta_ = 1.4 × 10^−20^) higher mean BMI per copy of the alternate (A) allele (Minster et al., 2016). A meta-analysis of Polynesians living in New Zealand showed an estimated 1.34 kg/m^2^ (*p*_meta_ = 1.60 × 10^−5^) higher mean BMI per copy of the alternate allele (Krishnan et al., 2018).

BMI is often used as a measure of body adiposity independent of height, although BMI is not consistently a stature-independent measure of adiposity, especially in children and adolescents (Benn, 1971; Cole, 1986; J. C. Wells, 2014; J. Wells & Cole, 2002). Thus, we hypothesized that rs373863828, in addition to having a strong association with BMI, could also be associated with height. We sought to characterize this potential relationship in Samoan adults and children.

## Methods

### Study Sample and Genotyping

We used three cohorts in this study: an adult discovery cohort of 3,077 Samoans, an adult replication cohort of 2,103 Samoans and American Samoans, and a cohort of 409 Samoan children (Table 1). For all cohorts, height was recorded using a portable GPM anthropometer (Pfister Imports, New York, NY) and rs373863828 was genotyped using a TaqMan real-time PCR assay (Applied Biosystems), as previously described (Minster et al., 2016). These studies were approved by the Health Research Committee of the Samoan Ministry of Health, the American Samoan Department of Health Institutional Review Board (IRB; for Samoa and American Samoa-based studies respectively) and the Brown University IRB. All participants gave written informed consent.

**Table 1.**
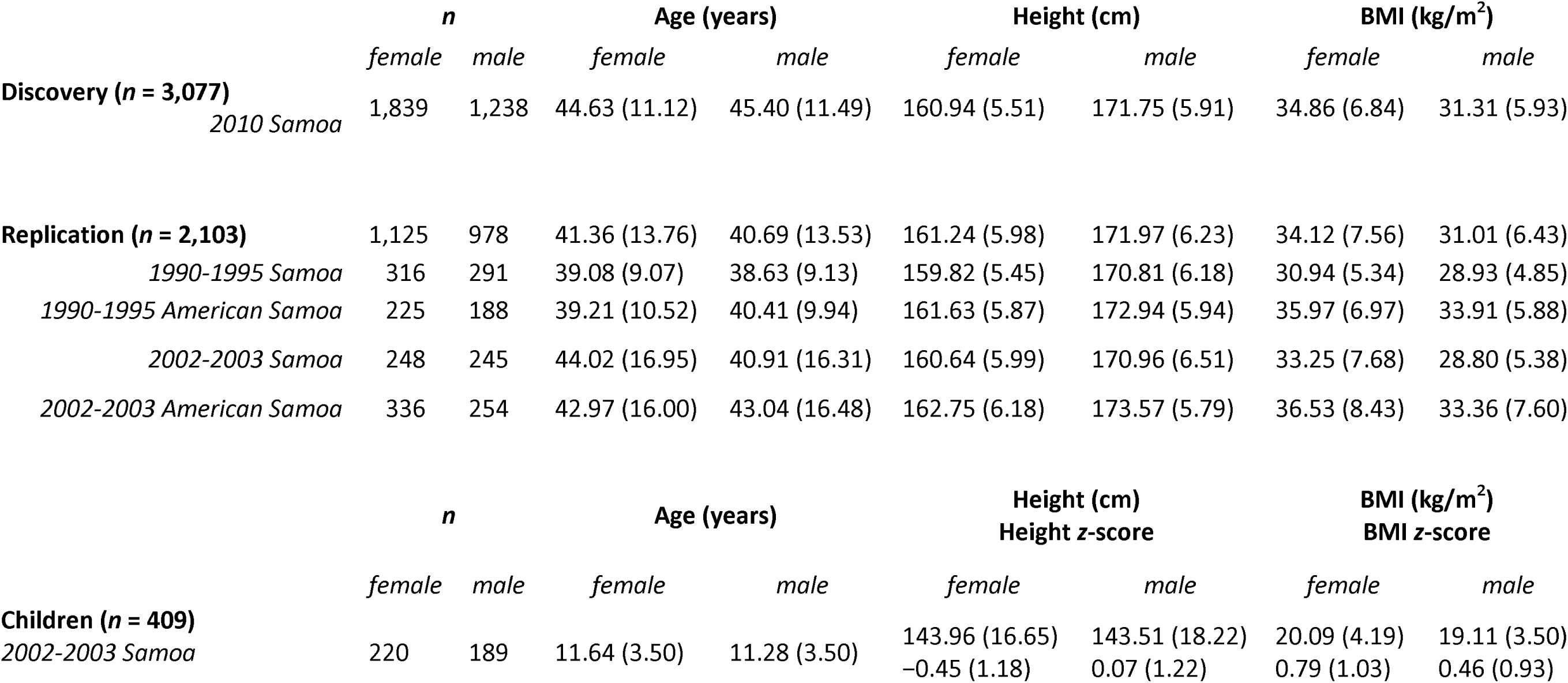
Sample characteristics for males and females in each cohort. Mean (SD) is given for age, height, and BMI for adults and children. Additionally in children, mean (SD) is given for age- and sex-specific height and BMI z-scores based on World Health Organization criteria.

#### Discovery Cohort

The discovery cohort of 3,077 adults of Samoan ancestry, 23–70 years old, was drawn from a population-based sample recruited from 33 villages located across the two major islands of the Independent State of Samoa (hereinafter Samoa) (Table 1). The sample selection, data collection methods, phenotyping and extensive quality control using genome-wide genotyping data has been reported previously (Hawley et al., 2014; Minster et al., 2016).

#### Replication and Children Cohorts

The replication cohort totaling 2,103 adults was taken from two studies from Samoa and the United States Territory of American Samoa (hereinafter American Samoa) (Table 1). The first sample totaling 1,020 participants was derived from a longitudinal study of adiposity and cardiovascular disease risk factors among adults from American Samoa and Samoa recruited between 1990 and 1995, using stature as recorded at baseline. Detailed descriptions of the sampling, recruitment, and phenotyping have been reported previously (Chin-Hong & McGarvey, 1996; Galanis, McGarvey, Sobal, Bausserman, & Levinson, 1995; McGarvey, Levinson, Bausser-Man, Galanis, & Hornick, 1993). Due to lack of genome-wide marker data on these samples, we were unable to infer relatedness, and so the participants were treated as unrelated in the analyses.

The second sample of 1,083 adults from American Samoa and Samoa, recruited in 2002–2003, was drawn from an extended family-based genetic linkage study of cardiometabolic traits (Åberg, Dai, Sun, et al., 2009; Åberg et al., 2008; Åberg, Dai, Viali, et al., 2009). Probands and relatives were unselected for obesity or related phenotypes, and all individuals self-reported Samoan ancestry. The recruitment process, criteria used for inclusion in this study, and phenotyping have been described in detail previously (Dai et al., 2007, 2008). The child cohort comprises 409 Samoan children, 5–18 years old, who are members of the recruited families. Relatedness was derived from known family structures which have been verified to be consistent with relatedness estimates derived using genome-wide microsatellite markers (Dai et al., 2007, 2008).

### Statistical Analysis

The association between rs373863828 and height was assessed separately for adults and children to account for differences in height during maturation and adulthood. In adults, the association was tested for separately in the discovery cohort of 3,077 adults and in the replication cohort of 2,103 adults.

In the discovery cohort, height residuals were calculated adjusting for the fixed effects age, age^2^, sex, age × sex interaction, and age^2^ × sex interaction. An empirical kinship matrix was generated with KING (Manichaikul et al., 2010) and PC-Relate (Conomos et al., 2019) to distinguish recent and distant relatedness in the sample using markers from the Genome-Wide Human SNP 6.0 array (Affymetrix). The percent of total variation in height and BMI was estimated using the coefficient of determination (R^2^) from a simple linear regression of each phenotype as predicted by an additive effect of rs373863828.

In the replication cohort, height residuals were calculated adjusting for the fixed effects of study (1994-95 or 2002-03 recruitment), polity (American Samoan or Samoa), age, age^2^, sex, age × sex, and age^2^ × sex. Kinship was estimated from pedigree information with the kinship2 package in R (Therneau & Sinnwell, 2014).

In children, the association was measured in the 409 children from the replication cohort. First, age- and sex-specific z-scores of height were calculated for children based on World Health Organization (WHO) References 2007 (de Onis et al., 2007) (Table 1). The resulting height z-scores were adjusted for the effects of age, age^2^, sex, age × sex, and age^2^ × sex. Kinship was estimated from pedigree information with the kinship2 package (Therneau & Sinnwell, 2014).

Within each cohort, age was centered to the overall mean for that cohort to avoid collinearity among the effects. Additionally, within each cohort, height (or height *z*-score) residuals were examined, and no evidence of non-normality was found (results not shown). These residuals were then used to assess the relationship between rs373863828 and height via linear mixed models adjusting for kinship as a random effect using the lmekin function in R (Therneau, 2018). The resulting effects from the two analyses in adults were combined using an inverse-variance effect-based fixed-effect meta-analysis as implemented in the rmeta package (Lumley, 2018).

Sex-specific effects of rs373863828 on height were also examined through a test of statistical interaction between sex and rs373863828 genotype in each cohort.

## Results

Association between rs373863828 and height was examined using linear mixed models in a discovery cohort of 3,077 adults and in a replication cohort of 2,103 adults, adjusting in the replication cohort analyses for the average effect of polity and study to account for the observed differences. Each cohort demonstrated significant association between the A allele of rs373863828 and taller average height (β_discovery_ = 0.60; 95% CI [0.30 - 0.91]; *p*_discovery_ = 1.00 × 10^−4^; β_replication_ = 1.06; 95% CI [0.65 −1.46]; *p*_replication_ = 2.80 × 10^−7^; Figure 1). Fixed-effects meta-analysis yielded a weighted effect estimate of 0.77 cm taller average height per copy of the A minor allele (95% CI [0.53 - 1.01]; *p*_meta_ = 5.56 × 10^−10^). The Cochran test for heterogeneity of effects was not significant (*p* = 0.08).

**Figure 1.**
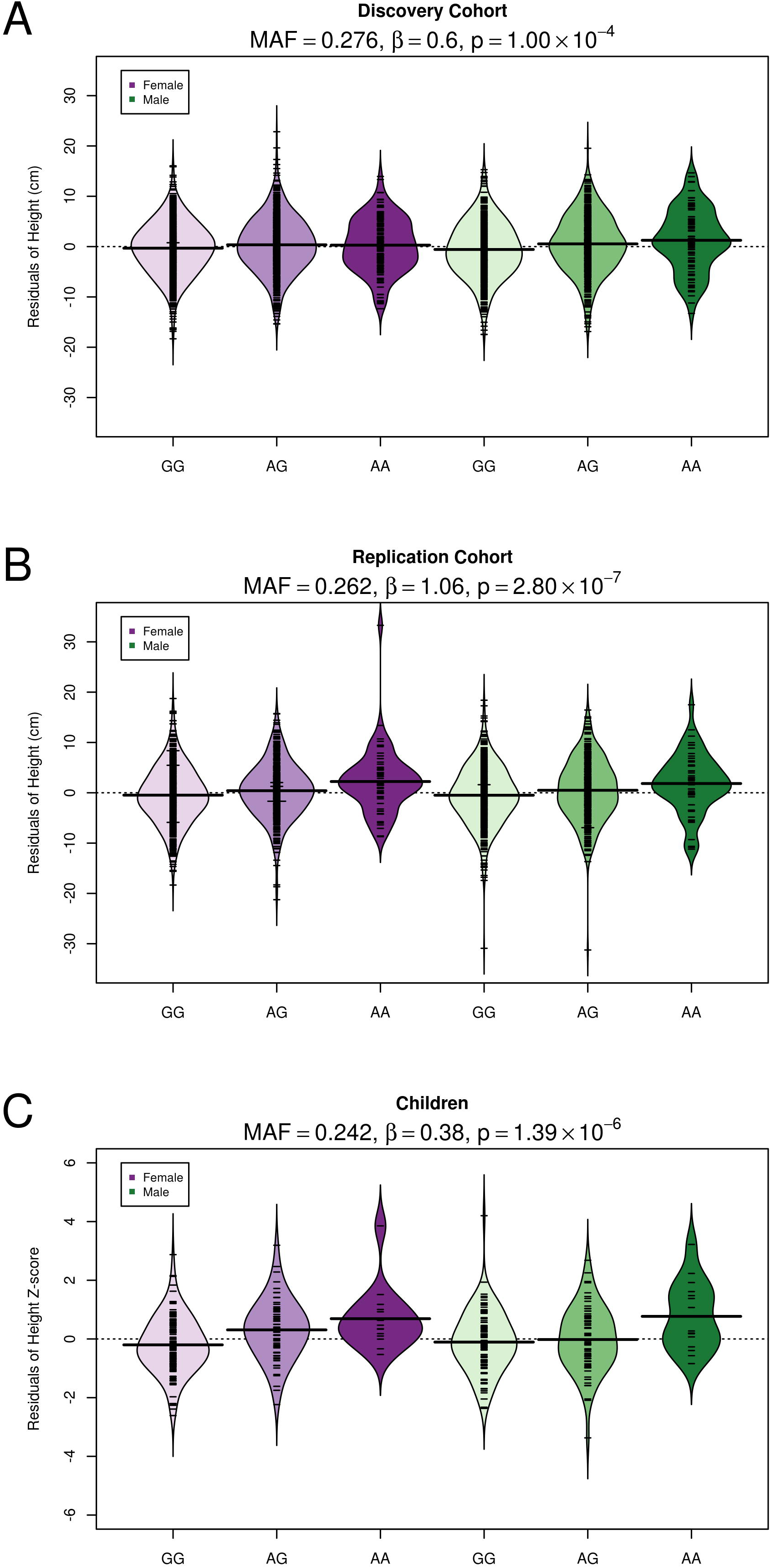
Beanplot displaying height residuals by rs373863828 genotype (GG, GA, AA) across sex and cohort. (A) Height residuals (cm) for the discovery cohort were adjusted for age, age^2^, sex, age × sex, and age^2^ × sex. (B) Height residuals (cm) for the replication cohort were adjusted for recruitment year, polity, age, age^2^, sex, age × sex, and age^2^ × sex. (C) Age- and sex-specific height z-scores based on World Health Organization criteria were adjusted additionally for age, age^2^, sex, age × sex, and age^2^ × sex. In each plot, individual observations are indicated with tick mark and the mean height residuals within each stratum is given by the black horizontal bar. Additionally, the minor allele frequency (MAF) of rs373863828, as well as the effect estimates (β) and p values from a linear mixed model testing the additive association between rs373863828 and height residuals are given within each cohort.

Notably, 0.18% of the total variation in unadjusted height was explained by an additive effect of rs373863828, compared to 1.95% of the total variation in unadjusted BMI.

The association between rs373863828 and taller stature was also observed in 409 children, with an effect estimate of 0.38 taller average height *z*-score per copy of the A allele (95% CI [0.23 - 0.53]; p = 1.39 × 10^−6^; Figure 1).

We also examined whether the effect of rs373863828 on average height differed for males and females in each cohort by testing for a statistical interaction between sex and genotype. We did not observe a significant interaction effect (β_discovery_ = 0.59, *p*_discovery_ = 0.04; β_replication_ = −0.05, *p*_replication_ = 0.09, β_meta_ = 0.38, *p*_meta_ = 0.11).

## Discussion

We have established an association of the minor allele (A) of rs373863828 with greater mean height in the Samoan population. This association confers an estimated taller mean height of 0.77 cm per copy of the minor allele in adults and an estimated 0.38 standard deviations taller height z-score in children, irrespective of sex. This observed effect was seen across two cohorts of adults recruited at three different time periods between 1990 and 2010.

A similar association between rs373863828 and height was observed in a cohort of Māori and Pacific adults from New Zealand (β = 1.25 cm; p = 3.90 × 10^−6^) (T. R. Merriman, personal communication, April 15, 2019). Additionally, association between rs373863828 and early childhood growth between ages 2 and 4 years old has been previously reported in a separate cohort including self-identified Māori and Pacific from New Zealand (Berry et al., 2018b, 2018a; Major et al., 2018).

This association, of rs373863828 with greater average height, is consistent in both adults ages 23–70 and children ages 5–18 years old, suggesting that the effect of rs373863828 on height may occur early in life. However, it is important to note that these data are cross-sectional and unadjusted for pubertal status, so future longitudinal studies will be required to determine if the effect of this variant on height can be attributed to overall growth trajectory or increased growth during a particular developmentally crucial period.

In addition to being one of the largest effect sizes reported for a common variant (β = 0.77, MAF = 0.259) on height, rs373863828 is a Pacific Island-specific polymorphism. Combined with previous association studies connecting rs373863828 with greater average adiposity and lower risk of diabetes, these results suggest that rs373863828 may operate broadly as a body size variant, affecting overall growth as opposed to a singular growth dimension. It is worth noting that this variant explains more of the variation in BMI than height (1.95% versus 0.18% respectively). However, the function of *CREBRF* is minimally characterized, so the mechanisms by which rs373863828 affects these phenotypes are currently unclear. Individual studies have indicated that this transcription factor impacts glucocorticoid signaling, autophagy, unfolded protein response, and starvation resistance, as well as murine decidualization, embryo implantation, and maternal behavior (Audas, Li, Liang, & Lu, 2008; Li et al., 2016; Martyn et al., 2012; Minster et al., 2016; Xue et al., 2016; Yang et al., 2013). Future studies should aim to refine the effect of this variation on the related phenotypes of height, weight, and BMI in other Pacific Island populations, as well as other anthropometric traits. Functional studies will be crucial to determining the function and targets of the *CREBRF* gene.

## Acknowledgements

The authors would like to thank the Samoan participants of the study, local village authorities, and the many Samoan and other field workers over the years. We acknowledge the Samoan Ministry of Health, the Samoa Bureau of Statistics, and the American Samoan Department of Health for their support of this research. We give particular thanks to two research assistants, Melania Selu and Vaimoana Lupematasila, who contributed to the 2010 recruitment and continue to assist us in our work in Samoa. This work was funded by the National Institute of Health grants R01-HL093093 (STM), R01-HL133040 (RLM), R01-AG09375 (STM), R01-HL52611(MI Kamboh), R01-DK59642 (STM), and R01-DK55406 (RD). Genotyping was performed in the Core Genotyping Laboratory at the University of Cincinnati, funded by National Institutes of Health grant P30-ES006096 (SM Ho).

## Author Contributions

JCC and SLR performed the statistical analysis, created the figures and tables, and wrote the manuscript, with guidance from RLM, STM, NLH, and DEW. EMR contributed to the statistical analysis under the guidance of RLM. NLH led the field work data collection and phenotype analyses with guidance from STM. GS led and directed genotyping experiments (using the Affymetrix 6.0 chip) and assay development for validation and replication (using the TaqMan platform) with guidance from RD. MSR facilitated fieldwork in Samoa. TN contributed to the discussion of the public health implications of the findings. All authors contributed to this work, discussed the results, and critically reviewed and revised the manuscript.

## Conflict of Interest Statement

DEW, NLH, RD, STM, and RLM are co-inventors on the United States Patent Application 20180245155, “Compositions and Methods for Identifying Genetic Predisposition to Obesity and for Enhancing Adipogenesis”.

